# Topographical similarity of cortical thickness represents generalized anxiety symptoms in adolescence

**DOI:** 10.1101/2023.03.09.531836

**Authors:** Chaebin Yoo, M. Justin Kim

## Abstract

Generalized anxiety disorder (GAD) is a common condition characterized by excessive, uncontrollable worry. Despite increasing efforts to identify the neural underpinnings of GAD, neuroimaging research using cortical thickness have yielded largely inconsistent results. To address this, we adopted an inter-subject representational similarity analysis framework and utilized a sample of 120 adolescents (13 to 18 years of age) from the Healthy Brain Network dataset. We found greater topographical resemblance among participants with heightened generalized anxiety symptoms in the left caudal anterior cingulate and pericalcarine cortex. Such associations were not observed when including a group of younger participants (11 to 12 years of age), highlighting the importance of age range selection when considering the link between cortical thickness and anxiety. Our findings reveal a novel cortical thickness topography that represents generalized anxiety in adolescents, which is embedded within the shared geometries between generalized anxiety symptoms and cortical thickness.

## Introduction

Generalized anxiety disorder (GAD) is characterized by excessive and uncontrollable worry, combined with symptoms such as restlessness, irritability, and bodily tension (VandenBos, 2007). Along with its high prevalence rate, GAD is known to increase risk for other psychological problems, showing high comorbidity with depressive and other anxiety disorders - exhibiting nearly 56-75% of comorbidity rates in youth (Masi et al., 2004). Of note, understanding early-onset GAD is of particular importance due to its potentially damaging consequences later in life. Indeed, a longitudinal study revealed that early GAD led to poor health, financial, and interpersonal outcomes later in young adulthood (Copeland et al., 2014). Considering such long-lasting negative influences, human neuroimaging studies have continuously searched for potential neuroanatomical markers of GAD (Zugman et al., 2022).

In an effort to identify more stable, trait-like underpinnings of mental disorders, several magnetic resonance imaging (MRI)-derived brain structural measures, including cortical thickness estimates, have been widely used in neuroimaging research. Cortical thickness is known to effectively reflect abnormal neuroanatomy, and to work as a potential biomarker offering information on disease progression, clinical diagnosis, and assessment (Hutton et al., 2008). Studies examining the relationship between cortical thickness and GAD, however, have previously yielded largely inconsistent results. Whereas the Enhancing Neuroimaging Genetics through Meta-Analysis (ENIGMA)-Anxiety Working Group study spanning 28 research sites and over 4,000 participants reported no linear relationship between GAD and cortical thickness (Harrewijn et al., 2021), other studies have suggested the contributions of the frontal (Abdallah et al., 2012; Cha et al., 2014; Strawn et al., 2014; Molent et al., 2018, Carnevali et al., 2019, Maggioni et al., 2019), parietal (Abdallah et al., 2012), temporal (Abdallah et al., 2012; Strawn et al., 2014), occipital (Abdallah et al., 2012; Strawn et al., 2014), and cingulate (Carnevali et al., 2019) cortices to GAD symptoms. Notably, there have also been contradictory findings where the orbitofrontal cortices of GAD patients have been reported to be thinner in some studies (Carnevali et al., 2019; Maggioni et al., 2019) and thicker in another (Abdallah et al., 2012). Although these inconsistencies may partly be due to differences in definitions and methodological approaches (Sobral et al, 2022), they may further imply that the cortical thickness–GAD relationship could be better captured through methods that examine beyond simple linear associations.

To address this, we utilized an inter-subject representational similarity analysis (IS-RSA) (Finn et al., 2020) framework to examine the multivariate patterns of similarity in vertex-wise cortical thickness associated with generalized anxiety symptoms. In IS-RSA, the pairwise geometry of brain data is compared with the pairwise geometry of behavioral data, using second-order statistics instead of directly associating two physically distinct quantities (Kriegeskorte & Kievit, 2013). Aiming to detect an “generalized anxiety archetype” of cortical thickness topography (i.e., vertex-wise arrangement of cortical thickness), we adopted the Anna Karenina (AnnaK) model (Finn et al., 2020) that assumes greater similarity among participants with higher anxiety scores. In other words, we sought to identify shared neural features representative of generalized anxiety in adolescents.

## Methods

### Participants

All participant data were obtained from the Healthy Brain Network (HBN) Biobank shared by the Child Mind Institute, an open dataset collected from a community sample of children and adolescents (Alexander et al., 2017). The present study utilized data from Release 7 to 10 scanned from two sites, the Rutgers University Brain Imaging Center (RUBIC) (*n* = 34) and the CitiGroup Cornell Brain Imaging Center (CBIC) (*n* = 94). Based on a prior study reporting distinct properties in cortical thickness between early and mid-adolescence divided with a boundary of age 13 (Mills et al., 2021), participants within the age range of 13 to 18 were included in the present study. Among the participants who have finished required questionnaires, those with T1 image preprocessing errors (*n* = 8) were excluded after visual inspection leaving 120 participants for final analyses (80 males, mean age 15.47 ± 1.59 years). 107 participants were newly diagnosed at the time of data acquisition of at least 1 psychopathology, including ADHD (*n* = 45), anxiety disorders (*n* = 28), depressive disorders (*n* = 12), autism spectrum disorder (*n* = 5), intellectual disability (*n* = 5), specific learning disorder (*n* = 4), language disorder (*n* = 2), adjustment disorder (*n* = 1), alcohol use disorder (*n* = 1), borderline intellectual functioning (*n* = 1), conduct disorder (*n* = 1), obsessive-compulsive disorder (*n* = 1), and substance-induced bipolar disorder (*n* = 1).

### Image Acquisition

Structural MRI images were acquired on a Siemens 3T Tim Trio scanner (RUBIC) and a Siemens 3T Prisma scanner (CBIC) with the same parameters. T1-weighted images covering the whole brain were obtained with a spatial resolution of 0.8 × 0.8 × 0.8 mm^3^ (224 slices). Other parameters were as follows: repetition time (TR) = 2500 ms, inversion time (TI) = 1060 ms, echo time (TE) = 3.15 ms, field of view (FOV) = 256 × 256 mm^2^, flip angle = 8°. Further details on scan protocols can be found here http://fcon_1000.projects.nitrc.org/indi/cmi_healthy_brain_network/MRI%20Protocol.html.

### Image Preprocessing

Anatomical data was preprocessed as part of the fMRIprep (ver 21.0.2) (Esteban et al., 2019) preprocessing pipeline. T1-weighted images initially went through bias field correction and were skull-stripped via ANTs workflows (Tustison et al., 2010). Then, *recon-all* in FreeSurfer (ver 6.0.1) (Dale et al., 1999) was run based on previously calculated brain masks. In this process, steps including tissue classification, surface reconstruction, cortical parcellation, and estimation of various surface data including cortical thickness took place. Images parcellated based on the Desikan-Killiany atlas (Desikan et al., 2006) were used for the current study, resulting in 33 regions per hemisphere. Finally, surface-source vertex-wise cortical thickness data were resampled onto the fsaverage template and concatenated to readable files using the *mris_preproc* function in FreeSurfer.

### Generalized anxiety symptoms

Generalized anxiety symptoms were measured with the Screen for Children Anxiety Related Disorders (SCARED; Birmaher et al., 1999). SCARED is a self-report, 41-item questionnaire that assesses symptoms related to five anxiety disorders parallel to DSM-? classification. Only the generalized anxiety disorder (GAD) symptom scores were used for analyses in the present study. We note that although the suggested cutoff that may indicate GAD is the score of 9 in SCARED, actual diagnoses are not necessarily in accord with this criterion.

### Inter-Subject Representational Similarity Analysis (IS-RSA)

In order to examine a potential nonlinear relationship between cortical thickness and generalized anxiety symptoms, inter-subject representational similarity analysis (IS-RSA) was conducted (Figure 1).

**Figure 1.**
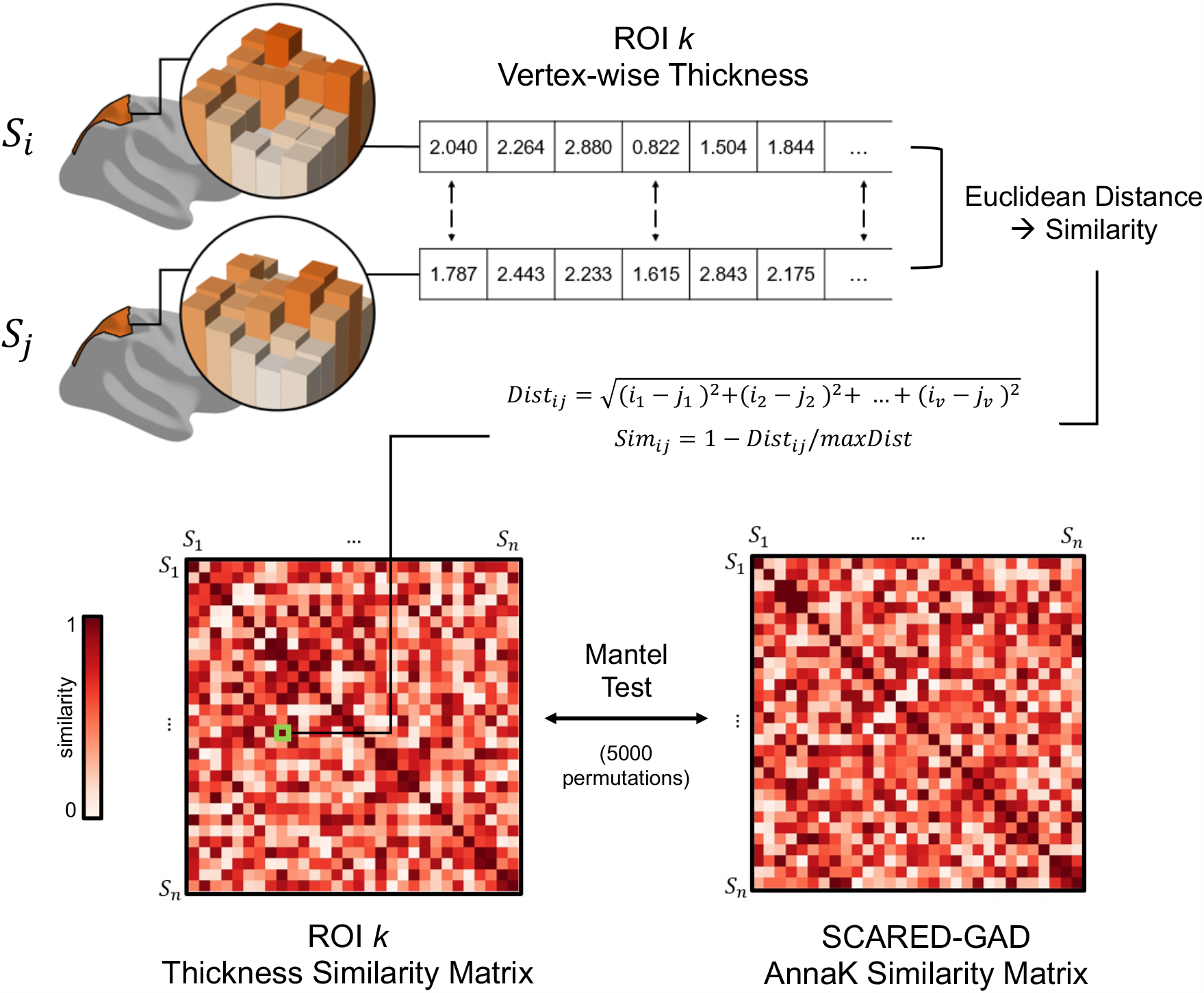
Inter-subject representational similarity analysis of cortical thickness. For each ROI, all participants’ vertex-wise cortical thickness indices were derived in vector form. Pairwise Euclidean distances of the vectors were computed, and converted to similarity indices, resulting in a 120 × 120 ROI thickness similarity matrix. This matrix was then compared with the generalized anxiety similarity matrix through permutation-based Mantel test. This process was repeated for all 66 ROIs defined by the Desikan-Killiany atlas. Values in the figure were generated for illustrative purposes. *Abbreviation:* ROI, region of interest.

#### Cortical Thickness Similarity Matrix

For each parcellated region, a 120 × 120 (i.e., subject × subject) cortical thickness similarity matrix was derived. First, vertex-wise thickness values that correspond to the specific region were extracted in vector form for each participant. Pairwise Euclidean distances were calculated between those vectors, and then converted to similarity through dividing them by the maximum distance and subtracting the value from 1.

Resulting indices were finally expressed in a symmetrical similarity matrix.

#### Generalized Anxiety Similarity Matrix

Since we assumed that participants with higher anxiety scores would exhibit similar cortical thickness topography, the Anna Karenina (AnnaK) similarity framework (Finn et al., 2020) was adopted. Here, generalized anxiety similarity was estimated as the pairwise mean of participants’ ranks based on their anxiety scores. Participants with lower generalized anxiety scores would thus be given lower similarity indices, whereas participants with higher scores would be given higher similarity indices. A 120 × 120 generalized anxiety similarity matrix was derived as the result.

#### IS-RSA

Spearman correlations between each region’s cortical thickness similarity matrix and the generalized anxiety similarity matrix were calculated. In other words, the geometric patterns of the two matrices were compared. The correlation coefficient would thus reflect actual similarity in brain topography among participants who show higher anxiety scores (i.e., who are assumed to be more similar). Significance of the correlation coefficient was tested by running 5,000 permutation tests and comparing the value with the null distribution built based on shuffled matrices. Results were corrected for multiple comparisons based on false discovery rate (FDR; *q* < .05) (Benjamini & Hochberg, 1995).

## Results

### Vertex-wise IS-RSA

Among all 66 cortical regions, significant correlations were found between the generalized anxiety similarity matrix and the vertex-wise cortical thickness similarity matrices of the left caudal anterior cingulate cortex (ACC) (*r* = .212, *p* = .002) and pericalcarine cortex (*r* = .223, *p* = .003) (Figure 2, Table 1). In other words, participants with higher generalized anxiety scores exhibited similar vertex-wise topography of cortical thickness in these regions. No region was found significant in the right hemisphere after correcting for multiple comparisons.

**Figure 2.**
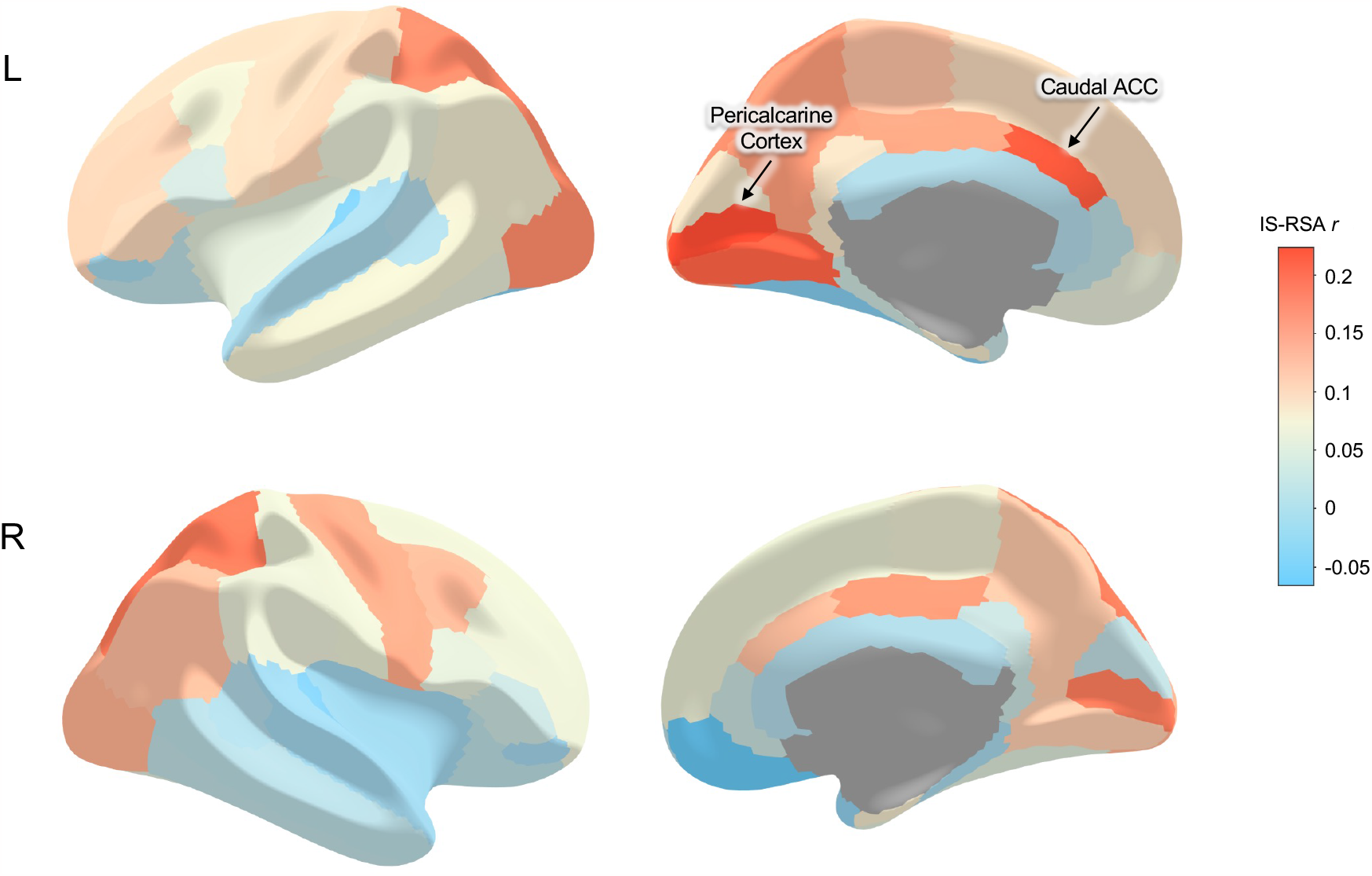
Results of the inter-subject representational similarity analyses shown from the lateral (left panel) and medial (right panel) view of the brain. Only the left caudal ACC and pericalcarine cortex were significant, indicating higher topographical resemblance in these regions among adolescents with heightened generalized anxiety. *Abbreviations:* ACC, anterior cingulate cortex; L, left; R, right.

**Table 1.** Regions showing high IS-RSA correlation coefficient in the main sample (*n* = 120). Results surviving the FDR-corrected threshold of *q* < .05 are marked with asterisks. *Abbreviations:* L, left; R, right.

| **Region** | ***r*** | ***p*** |
| --- | --- | --- |
| L caudal anterior cingulate | 0.212* | 0.002* |
| L pericalcarine | 0.223* | 0.003* |
| L lingual | 0.205 | 0.005 |
| R pericalcarine | 0.183 | 0.014 |
| R superior parietal | 0.185 | 0.022 |
| L superior parietal | 0.171 | 0.030 |
| L lateral occipital | 0.170 | 0.034 |
| R posterior cingulate | 0.159 | 0.042 |
| L posterior cingulate | 0.148 | 0.048 |

### Control Analyses

Six sets of control analyses were conducted to rule out potential alternative explanations. First, when using the *mean* cortical thickness measures instead of the vertex-wise indices, the generalized anxiety similarity matrix and the cortical thickness similarity matrices were no longer correlated in any of the 66 cortical regions (all *p*s >.119). Second, age similarity was not correlated with the left caudal ACC thickness similarity (*r* = .030, *p* = .692), nor with the left pericalcarine thickness similarity (*r* = .090, *p* = .206). Sex similarity also showed no correlation with similarities in either region (left caudal ACC, *r* = -.039, *p* = .264; left pericalcarine, *r* = -.013, *p* = .679). Third, when using the parent reports of the SCARED-GAD scores, no significant correlation was found between any of the regional cortical thickness similarity matrices and the anxiety similarity matrix (all *p*s > .133). Fourth, the depression similarity matrix estimated based on the Mood and Feelings Questionnaire (MFQ) (Costello & Angold, 1988) scores was not significantly correlated with any of the regional cortical thickness similarity matrices after multiple comparisons correction. We note that the left pericalcarine cortex thickness similarity was correlated with depression similarity at an uncorrected threshold (*r* = .157, *p* = .026). Fifth, after eliminating participants diagnosed with GAD (*n* = 14) from the analysis, significant correlations were still found between the generalized anxiety similarity matrix and the cortical thickness similarity matrices of the left pericalcarine cortex (*r* = .231, *p* = .002), lingual gyrus (*r* = .232, *p* = .004), and caudal ACC (*r* = .225, *p* = .004). Sixth, after eliminating participants diagnosed with ADHD (*n* = 45), no region survived the FDR-corrected threshold. However, results showed a similar tendency with the full sample overall, with higher correlations in the left lingual gyrus (*r* = .269, *p* = .006), caudal ACC (*r* = .248, *p* = .006), pericalcarine cortex (*r* = .246, *p* = .009), paracentral lobule (*r* = .210, *p* = .030), bilateral lateral occipital cortex (left, *r* = .259, *p* = .011; right, *r* = .225, *p* = .024), and superior parietal cortex (left, *r* = .242, *p* = .011; right, *r* = .252, *p* = .011).

### Generalizability to Early Adolescence

To determine whether these results could be generalized to a sample of younger, wider age range, we ran additional analyses after adding 120 participants between ages 11 to 12. This left us with 240 participants spanning ages 11 to 18 (161 males, mean age = 13.78 ± 2.09 years).

Analyses yielded parietal- and cortical midline-highlighted results, with significant correlations between the generalized anxiety similarity matrix and the cortical thickness similarity matrices of the left superior parietal cortex (*r* = .174, *p* = .001), precuneus (*r* = .155, *p* = .005), pars triangularis (*r* = .151, *p* = .005), pericalcarine cortex (*r* = .151, *p* = .005), postcentral gyrus (*r* = .150, *p* = .010), medial orbitofrontal cortex (*r* = .146, *p* = .010), and lateral occipital cortex (*r* = .146, *p* = .011). However, cortical thickness similarity matrices of these regions also showed significant correlations with the age similarity matrix (all *p*s < .007), failing to rule out the potential possibility of age driving the topographical similarity. Age similarity in this sample was also significantly correlated with similarity in generalized anxiety (*r* = .235, *p* = .0003), thus leaving the relationship between anxiety and cortex thickness uncertain in the wider, younger age range. When the analyses were restricted to 11-12-year-olds (*n* = 120), generalized anxiety similarity was not correlated with any of the regional cortical thickness similarity (all *p*s > .06).

## Discussion

Here, we utilized vertex-wise cortical thickness data of 120 adolescents to explore its potential nonlinear relationship with generalized anxiety symptom severity. Data-driven IS-RSA analyses revealed significant correlations between generalized anxiety similarity and cortical thickness similarities of the left caudal ACC and pericalcarine cortex, indicating greater topographical resemblance in these regions among more anxious adolescents. Further analyses revealed that age, sex, neurodevelopmental ADHD diagnosis, and GAD diagnosis did not account for the observed effects. Moreover, the present findings were specific to self-reported but not to parent-reported generalized anxiety scores. When expanding the sample age range to include 11- and 12-year-olds, additional regions in the parietal lobe were significantly associated with generalized anxiety symptoms; however, their regional thickness similarities also showed high correlations with age similarity. These findings reveal a novel brain-anxiety association in adolescence, which is embedded within the shared geometries between generalized anxiety symptoms and cortical thickness.

Adolescents with heightened generalized anxiety symptoms shared similar cortical thickness topography in the left caudal ACC with each other, more so than those with lower anxiety. The ACC, one of the most frequently targeted brain regions in GAD research (Kim & Kim, 2021), is generally understood to be important for generating worry – a defining characteristic of GAD (Comer et al., 2004). In particular, the caudal ACC is known to be associated with fear expression, especially in regulating heart rate and skin conductance response (Heilbronner & Hayden, 2016; Milad et al., 2007). This is in line with the fact that GAD exhibits physiological arousal as one of its core symptoms, which may lead to perceptual distortions of their bodily states (Hoehn-Saric & McLeod, 2000). Using cortical thickness metrics, several studies have previously reported linear relationships between anxiety and ACC thickness. However, the results were inconsistent, with one study reporting thicker ACC for those with higher anxiety scores (Donzuso et al., 2014) while another noting thinner ACC in GAD patients (Carnevali et al., 2019). The current findings may aid in resolving the apparent inconsistencies by revealing a nonlinear relationship between generalized anxiety symptoms and topographical configurations of the ACC.

Another region that exhibited a similar effect was the pericalcarine cortex. The pericalcarine cortex, where the primary visual cortex is located, has been understood to be implicated in early threat perception and evaluation through fear conditioning (Miskovic & Keil, 2012; Wieser & Keil, 2020). Although the amygdala is usually thought to be at the core of threat learning, sensory cortices could also be recruited in early threat processing and may eventually take over threat learning via their long-term plasticity (Wieser & Keil, 2020). Interestingly, pericalcarine cortex thickness is associated with anxiety treatment outcome in children and adolescents (Gold et al., 2017) along with the tendency to engage in risky behaviors (Miglin et al., 2019). These findings imply that abnormalities in this region may lead to disrupted appraisal of threat, and that this disruption could be persistent. We can speculate, then, certain topographical characteristics of the pericalcarine cortex that are shared among teens with heightened generalized anxiety symptoms may reflect a similar, oversensitive tendency when evaluating encountered stimuli.

We attempted to generalize the current findings to early teens by including 11- and 12-year-olds in our analyses but found that the effects of age could not be ruled out from interpreting the results. Although there were several parietal and cortical midline regions whose thickness was significantly related to generalized anxiety scores, they also exhibited higher similarity among older adolescents. In fact, the majority of cortical thickness similarity across the whole brain was explained by age when the study sample was expanded to include participants with 11-18 years of age. On the other hand, age similarity showed no correlation with cortical thickness similarity in most regions, including those previously related to anxiety, when the study sample was restricted to 13-18 years of age. Thus, it can be inferred that vertex-wise cortical thickness topography generally shows convergence approximately post-age 13, which is broadly in line with the previous study reporting decreased variability in mean cortical thickness during mid-adolescence compared to an earlier period (Mills et al., 2021). As it has been pointed out before (Molent et al., 2018), age may be a crucial determinant of the results especially in studies employing cortical thickness metrics. The result of the present work suggests that the participant age range may need to be carefully considered, especially when studying youth.

Our findings further support the ongoing call for the reconsideration of the current classification of psychiatric disorders and their diagnosis criteria. As it has been pointed out (Fava & Mangelli, 2001), the current diagnostic criteria such as the DSM are mostly focused on determining whether one’s syndromes are deemed “clinical” or not, based on a cutoff. Many studies have thus been comparing clinical groups with “healthy” controls, disregarding subclinical symptoms in the process. However, it is important to note that including or excluding patients who were categorically diagnosed of GAD from IS-RSA had no bearing on the present findings. That is, the topographical characteristics of cortical thickness reflecting higher generalized anxiety dimension were identified regardless of GAD diagnoses. This implies that, at least when considering cortical thickness measures, there may not be a clear cutoff that distinguishes clinical and nonclinical groups. In addition, it is also noteworthy that categorical boundaries suggested by the current diagnostic criteria may not be as conclusive as previously assumed (Casey et al., 2013). Indeed, our findings showing overlapping cortical thickness similarities (albeit at an uncorrected level) associated with both generalized anxiety and depression symptoms are in line with alternative frameworks such as the Research Domain Criteria (RDoC), whereby psychopathology is explained with domains of basic neurobehavioral functioning (Sanislow et al., 2010).

Additionally, cortical thickness topography was no longer associated with generalized anxiety symptoms when parent-reported scores were used instead of self-reported scores. There have been questions regarding the substitutability of parent reports for children’s experiences, especially with studies reporting lower parent-child agreement for non-observable functioning such as emotion (Eiser & Mores, 2001). The present results provide partial support to this assertion, in that extracting topographical characteristics associated with generalized anxiety symptoms requires self-reported, but not parent-reported assessments.

There are a number of limitations that could be addressed in future research. First, the HBN dataset used for the present study was acquired from children and adolescents with psychiatric concerns. In fact, the majority of participants in the current sample was diagnosed with at least one psychopathology in the course of data collection. Therefore, the generalizability of the results to a more representative population would need to be tested. Also, assessment of generalized anxiety symptoms relied on either self-reported or parent-reported scores. Utilizing a more objective index of generalized anxiety (e.g., behavioral observation) would help to mitigate the shortcomings of self-reported measures. Finally, while we analyzed a sample of younger population to examine the generalizability of our results, it is yet to be confirmed whether the present findings can also account for older populations. A previous study examining brain developmental patterns (Mills et al., 2021) reported different trajectories in cortical thickness change not only between early teens and mid-adolescence, but also between mid-adolescence and adulthood. Thus, future studies would benefit from employing larger study samples with a wider age range that includes adults.

These limitations notwithstanding, the present study contributes to the relevant literature by identifying cortical regions – caudal ACC and pericalcarine cortex – that exhibit topographical resemblance among adolescents with heightened generalized anxiety symptoms. Our study also highlights the importance of age range selection when exploring the link between affect and brain structures, in that vertex-wise cortex thickness topography likely reflects brain maturation patterns in early teens. Our findings complement previous research in discovering trait-like brain structural underpinnings of adolescence anxiety disorders and may potentially serve as a neuroimaging-based diagnostic aid.

## Data Availability

The HBN dataset (http://fcon_1000.projects.nitrc.org/indi/cmi_healthy_brain_network/) is publicly available.

This research was supported by the National Research Foundation of Korea (NRF-2021R1F1A1045988). We thank the original authors of the HBN dataset for their generosity in making it available for use.

## Author Contributions

C.Y. and M.J.K. developed the study concept; C.Y. analyzed the data under the supervision of M.J.K; C.Y. and M.J.K. drafted the manuscript. All authors reviewed and approved the final manuscript for submission.

## Competing Interests

The authors declare that they have no conflict of interest.

## Notes

### Competing Interest Statement

The authors have declared no competing interest.

